# Morphoelastic modeling of pattern development in the petal epidermal cell cuticle

**DOI:** 10.1101/2022.10.30.514439

**Authors:** Carlos A. Lugo, Chiara Airoldi, Chao Chen, Alfred J. Crosby, Beverley J. Glover

## Abstract

We use the model system *Hibiscus trionum* as a vehicle to study the origin and propagation of surface nano-ridges in plant petal epidermal cells by tracking the development of the cell shape and the cuticle. In this system, the cuticle develops two distinct subdomains, (i) an uppermost layer which increases in thickness and in-plane extension and (ii) a substrate. We quantify the pattern formation and geometrical changes and then postulate a mechanical model assuming that the cuticle behaves as a growing bi-layer. The model is a quasi-static morpho-elastic system and it is numerically investigated in two and three dimensional settings, using different laws of film and substrate expansion and boundary conditions. We recreate several features of the observed developmental trajectories in petals. We establish the respective roles of the layers’ stiffness mismatch, the underlying cell-wall curvature, the cell in-plane expansion and the thickness growth rates of the layers in determining the observed pattern features, such as the variance observed in amplitude and wavelength. Our observations provide evidence which justify the growing bi-layer description, and provide valuable insights into why some systems develop surface patterns and others do not.

## 1 Introduction

The variety of topographic patterns found on the surface of plant petal and leaf epidermal cells has attracted the attention of researchers from disciplines ranging from developmental biology [1] [2] to material sciences [3]. These patterns have been shown to mediate several forms of interaction with the environment. For instance, it has been demonstrated [1, 4, 5] that patterned petals can act as optical and tactile cues for pollinators, providing some flowers with structural colour, and may also contribute to altered flower-pollinator adhesion or regulate hydrophobicity [6].

A striking example of pattern functionality is provided by the model system *Hibiscus trionum* (Fig. 1) in which the pattern consists of quasi-periodic nano ridges acting as a light diffracting grating array, conferring structural colour to the flower. The pattern forms on one of the two sub-domains of the petal’s adaxial surface which are defined by their epidermal cell shapes and which roughly coincide with the white and pigmented domains. The white domain consists of conical shaped cells that do not develop surface pattern. The rest of the petal is composed of cells with an elongated in-plane rectangular shape located mostly in the purple pigmented region of the tissue and where the cell surface develops a distinctive striated pattern.

**Figure 1:**
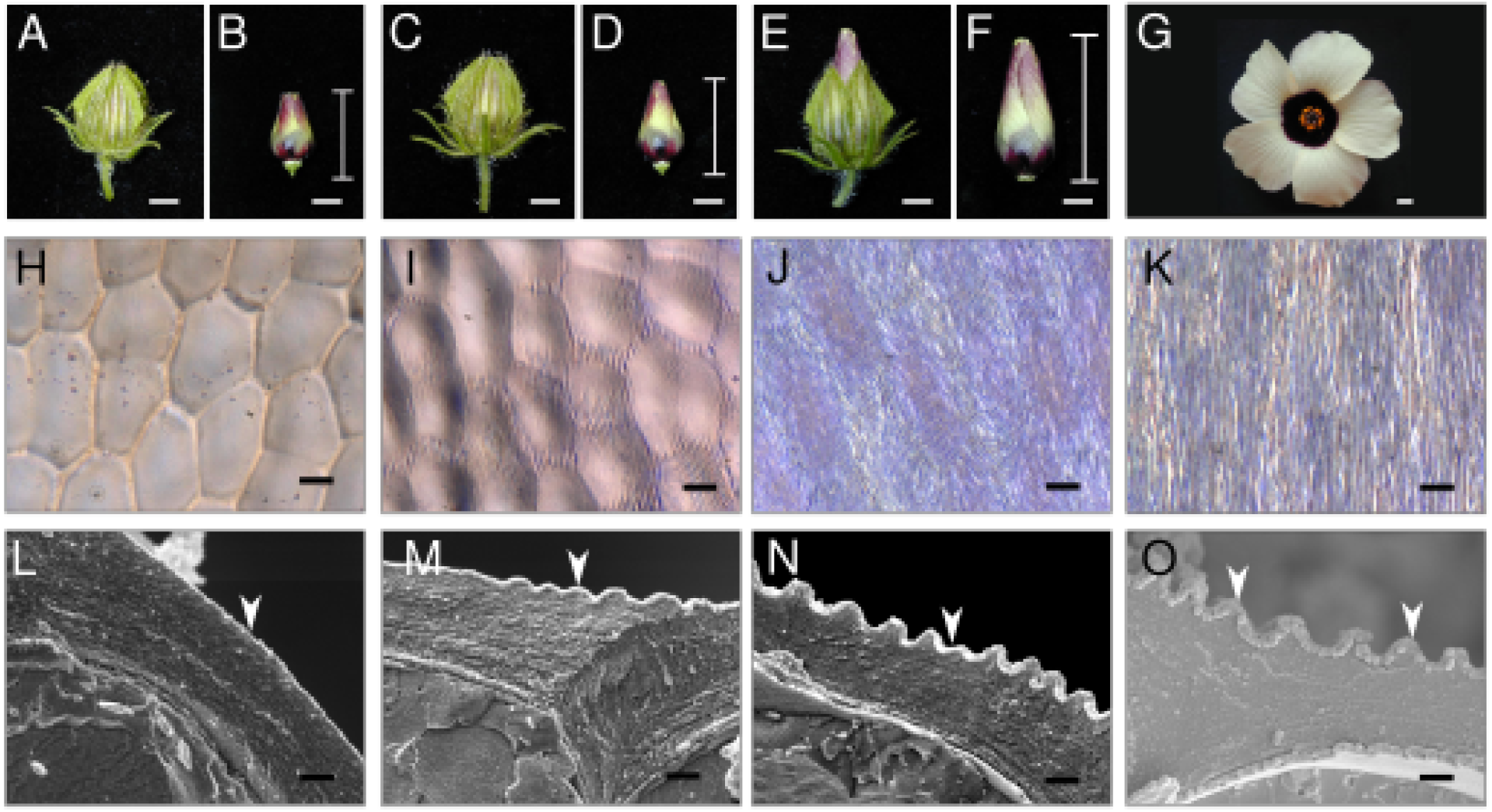
Development of *Hibiscus trionum* flower, epidermal cells, and surface morphologies. A-G Pictures of *Hibiscus trionum* buds (A–F) and an open flower (G), scale bars 1mm. Petals of the flower buds are covered by an epicalyx that has been removed in B,D,F. The bud size is referred to as the bottom to tip length after the epicalyx has been removed. H,I,J,K Keyence images of epidermal cell surfaces in the purple adaxial region of the petal at the stage shown above (scale bar 10 *μm*). L,M,N,O CryoSEM-fractures of *Hibiscus trionum* petal epidermal cells at the stage shown above (scale bars 1μm). A,B,H,L are pictures of a bud during the pre-striation stage (10 mm bud, measured as specified in the methods); C,D,I,M at the onset of striation (12.5mm-13mm bud); E,F,J,N are from a mature bud one day before flower opening (19-20mm bud). G,K,O pictures are from an open flower. The arrows in (L-O) indicate the development of the cuticle film.

The striated pattern aligns parallel to the major axis of the cells and forms on the cuticle, which is the outermost layer of the epidermis. Plant cuticles are extracellular polymer layers that act as a physical defence against a number of environmental agents. They comprise two chemically distinguishable sub-layers: a base layer that starts at the cell wall, which we refer to as the “cuticular substrate”, and an upper surface-most layer or “cuticle film” [7, 8, 9]. Such bi-layered structures play a significant role in systems that develop periodic surface deformations, as a result of a compression induced elastic instability. In the case of *Hibiscus trionum*, it has been shown that the cuticle behaves like a bi-layer composite, as the patterns can be induced by uniaxial stretching of the cuticles before the pattern occurs naturally [10]. The striated domain generates an iridescent reflectance spectrum which as reported in [3] is also different from that of a perfectly flat periodic grating and has served as inspiration in the design of new materials.

The origin of the surface deformation has been hypothesised to be mechanical in nature. Specifically, it may be due to a mechanical instability associated with cuticle overproduction. In [11] the authors assume the cuticle to be a homogeneous incompressible, mechanically isotropic layer, and propose that cuticle production is described by an adjustable parameter, which turns the in-plane stress components into tensile or compressive as the cuticle is anisotropically stretched due to cellular in-plane growth. This purely elastic formulation, however, does not address issues such as the existence of buckled configurations associated with the stress tensors proposed, the irreversible nature of the patterns or the critical nature of the elastic instability i.e., the existence of in-plane stress threshold values which need to be exceeded for the material to buckle. Nor does it provide any understanding of pattern features such as the amplitude or wavelength of striations and their relation to the material properties of the cuticle. A more comprehensive description of the mechanics behind patterned cuticles in petals is given in [12] by considering the cuticle as a bi-layered composite described by linear stress-strain constitutive relations and linking stress to in-plane cell and petal shape geometrical changes, which then determine the pattern orientation but without explicitly addressing the developmental aspects we discuss here.

A bilayer composite develops a surface periodic pattern or wrinkling/buckling instability in the upper surface if an applied in-plane stress in a given direction exceeds a threshold value [13, 14]. The resulting pattern is oriented in the perpendicular direction to the applied in-plane stress [15, 16, 17, 18, 19]. The critical threshold and pattern features depend on the elastic properties of the composite, its geometry and boundary conditions. For rectangular flat volumes, these dependencies are fairly general, regardless of the material model specifics, provided the elastic responses are isotropic [17, 15, 19], and are given by the scaling rules at the onset of the instability, namely, the wavelength:

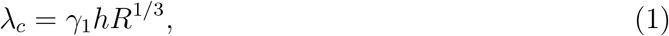

and the critical strain:

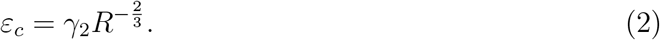

Where *h* is the film thickness, *R* the ratio of the elastic response modulus of the film and substrate and *γ*_1_ and *γ*_2_ are constants. Eqn 1 states that a thicker upper film induces a larger wavelength, whereas Eqn. 2 means that larger elastic mismatch between the film/substrate stiffness result in lower buckling thresholds. As described elsewhere [20, 17], these patterns emerge as trade-off states between the energetically favourable states of the slender film and the substrate under the same stress.

The universality of the buckling mechanism [13, 21, 22] means that the trends in Eqn. 1 and Eqn. 2 are followed by a large class of material models, regardless of the origin of the in-plane stress. For instance, periodic buckling patterns can be achieved by several settings involving uni-axial compression or unidirectional in-plane growth of the layers [23]. If in-plane stress acts on more than one direction, the possible instabilities which can be triggered range from checkerboards and hexagonal arrays to herringbones [19, 20].

In this work, we present results of a detailed developmental analysis of the cell geometry and pattern features in the *H. trionum* petal, which are then used to formulate morpho-mechanical models of the system [24]. As discussed below, this results in configurations of an irreversible nature and with deformations which are beyond the linear theory. Investigation of the models allowed us to elucidate the interplay of cuticle production, cell elongation and cell bulging in shaping the pattern and also permit us to pinpoint the locations where the pattern is triggered. Specifically, we explore the effects that different growth rates have on both reaching the buckling threshold and on the pattern features. Our results explain why some plant cuticles form surface patterns whereas others do not and clarify how this is related to cuticle synthesis and cuticle structural and mechanical properties [25]. Our results are also relevant in the development of bio-inspired engineered materials where some degree of tunability is a desired feature [3].

## Results

From a petal development perspective, it is not intuitive to understand the mechanism behind the source of stress which may induce a buckling instability. As we show next, quantitative measurements of changes of the cell geometry support the case for in-plane compressive stress induced by a larger expansion of the cuticle film in comparison to that of the substrate and underlying cells, in the direction orthogonal to the pattern ridges, to be the mechanism responsible for the origin of the buckling instability in H. *trionum*.

### Description of Hibiscus flower epidermis and its development

In *Hibiscus trionum* the pattern occurs in the uppermost layer on the external face of the epidermal cells, i.e. in the plant cuticle. The pattern develops from a flat surface in the early stages of petal development (Fig. 1 A,B,H,L). During the onset-stage striations appear first around the cell edges (Fig. 1 C,D,I,M). The ridges populate the entire cell surface at the mature bud stage (Fig. 1 E,F,J,N), assuming their final semi-ordered configuration in the open flower (Fig. 1 G,K,O).

The sequence in Fig. 1 (L-O) and additional cryo sections for before and after pattern formation (Fig. 2a (A-B)), show that the cuticle film layer is not distinguishable from the rest of the cuticle at an early stage of development, but appears very clearly as a distinct layer in the fully developed phenotype. The fully formed pattern image in Fig. 2a (B) shows features used to characterise the pattern, namely the amplitude *A*, wavelength λ and the cuticle film thickness *h*.

**Figure 2:**
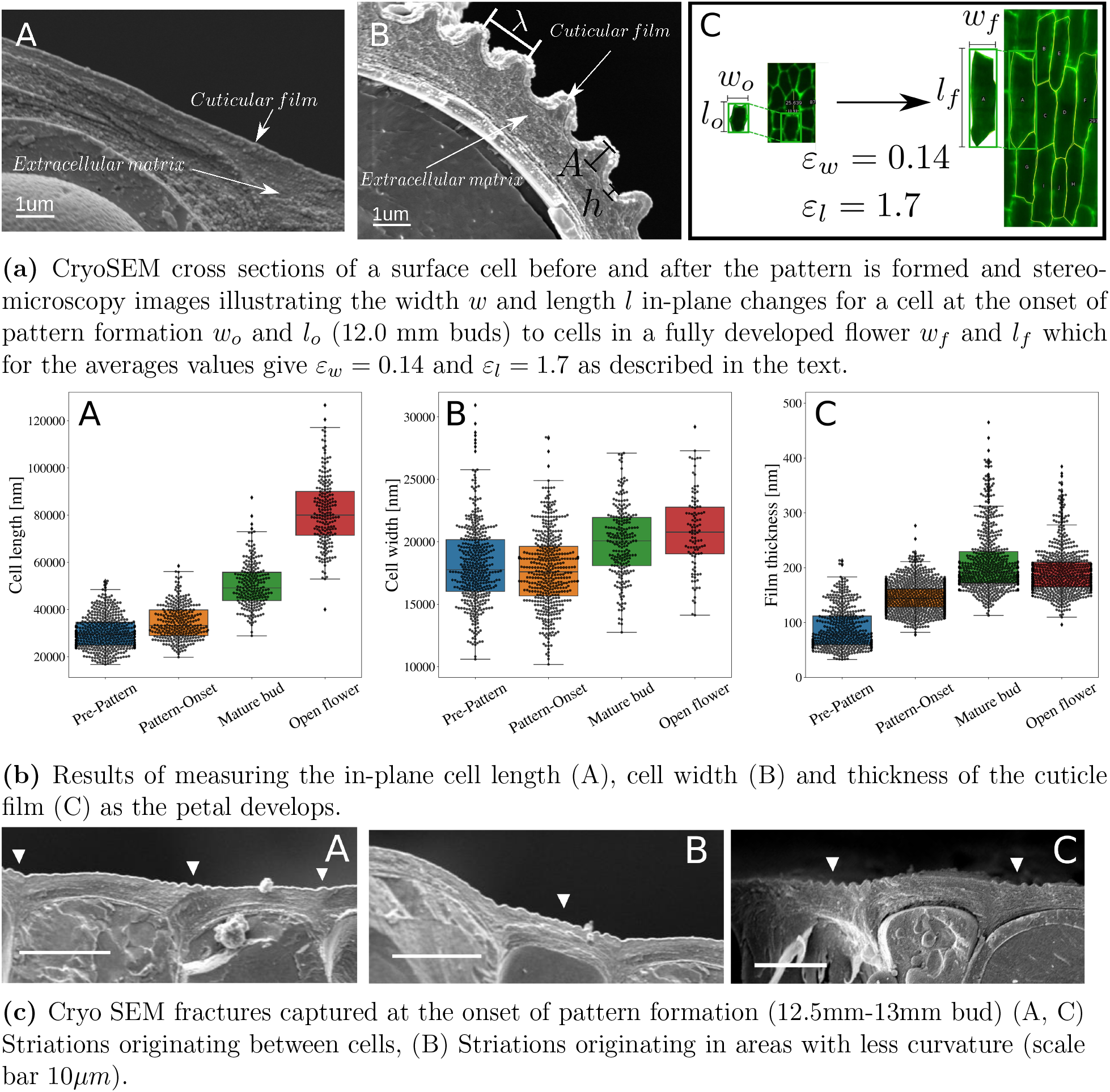
In-plane geometrical and morphological surface changes during development of *H. trionum* petal epidermal cells. (a) The cuticle film thickens and the cells elongate as the petal develops. (b) Box plots of the change in cell length (A), cell width (B) and (C) cuticle film thickness. (c) The pattern starts forming on flatter regions before developing across the entire domain which exhibits structural colour.

Both the in-plane width *w* and length *l* increase, except for a slight decrease in *w* before the striations start to form, where cell division is still more frequent than in-plane expansion. Fig. 2b shows measurements of these as well as for the thickness of the cuticle film. The changes 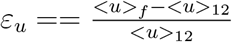 of the average in-plane cell width and length < *u* > (*u* ∈ {*w, l*}) with respect to the average cell widths and lengths for buds of 12 mm < *u* >_12_ obtained from the data are *ε_w_* ~ 0.14 and *ε_l_* ~ 1.7. Both show a net increase in both directions, most noticeable in the direction which grows parallel to the petal length, as shown in Fig. 1 (L-O). The thickness *h* of the cuticle film increases from a very thin layer to a value which remains almost constant during development. From the start of pattern formation until the flower opens, cell elongation is the main expansion mechanism responsible for the petal surface growth. The in-plane cell expansion and the cuticle thickening indicate that the volume of the cuticle layer is increasing, if such expansion is isotropic, then the cell expansion of the cells in a preferential (length) direction will induce an excess of cuticle in the perpendicular direction (width) effectively compressing the cuticle in that direction.

Fig. 2c shows several cross sections taken at the onset of pattern appearance. The pattern starts populating flatter sub-domains of the tissue such as the regions located on top of cell junctions, whereas for regions with larger curvature, pattern initiation is delayed by the bulging of the cell wall due to turgor pressure. This external domain deformation also contributes to a stress distribution as the regions with local positive signed curvature (concave) are effectively compressed, whereas those with negative signed curvature (convex) are under tension.

### Mechanics

We consider the cuticle as a bilayer volume consisting of an elastic film resting over a compliant substrate such as that shown in Fig. 3. The layers are characterised by their elastic properties. Assuming both the cuticle film and substrate cuticular layer to be mechanically isotropic, it is enough to specify the Lamé parameters of each layer to specify a constitutive relation between stress and deformation to model the passive responses.

**Figure 3:**
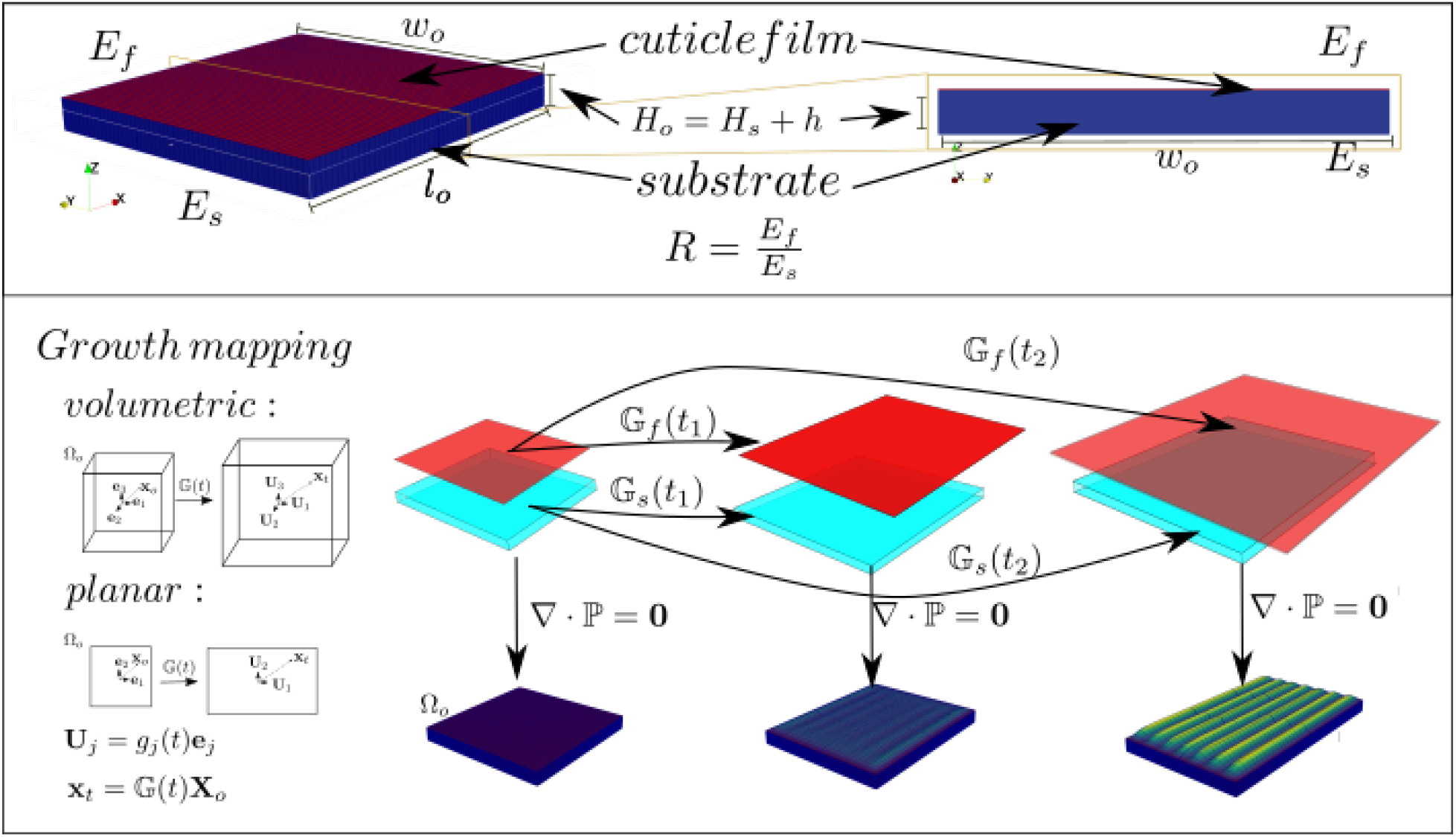
Cuticle bilayer model and the underlying geometry of growth induced stress. (Top) The reference configuration Ω_*o*_ consists of a volume of initial depth *H_o_* composed of two layers of initial thickess *H_s_* and *h* for the substrate and the cuticular film and the in-plane initial width and lengths *w_o_* and *l_o_*. These subdomains are characterised by their Young’s modulus *E_f_* and *E_s_*, which we encode in the parameter *R* = *E_f_/E_s_*. (Bottom) The geometry associated with the growth tensors 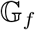 and 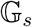 which map the reference configuration to “grown” configurations in terms of principal directions. These new configurations need to fulfill the imposed boundary conditions of the elastic problem which, as the figure shows, correspond to calculating the elastic response of accommodating a larger volume into a smaller one which is spanned by the underlying cell in-plane surface.

In this work we make use of hyperelasticity to describe the cuticles and specify the Lamé numbers in terms of the linear response modulus or Young’s modulus and Poisson’s ratio of each layer. If we consider the cuticle as a near-incompressible material across all the volume, then the latter *ν_s_* = *ν_f_* = *ν* ~ 0.5 where *f* and *s* stand for film and substrate. The linear response coefficients can be encoded in the stiffness mismatch ratio 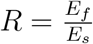.

A key difference between synthetic materials and living systems, such as petal surfaces, is that for living matter growth and remodelling are mostly due to internal processes rather than caused by the action of external agents such as an imposed displacement or an external force. For the case under consideration here, we assume that cuticle overproduction translates into a local increase of the volume occupied by every volume element composing the layers. The incorporation of growth into the mechanics framework or morpho-elasticity is based on the decomposition of the deformation gradient 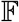 as a product of the elastic response 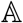 and a growth tensor 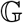.

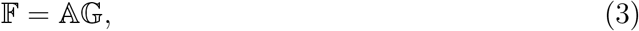

The decomposition Eqn. 3 allows us to solve an elastic problem provided we know the underlying growth rates of the solid with respect to a stress-free reference configuration Ω_o_. The complete formulation of the problem requires us to provide a constitutive equation for the material, boundary conditions and the fulfillment of the relevant balance laws, namely linear momentum, torque and mass. Assuming that the addition of mass into the system is such that the volumetric density remains approximately constant, then the stationary configuration is found by solving the elastic problem:

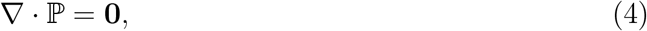

where 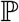 is the nominal stress tensor (see Methods). The components of the tensor 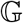 provide the link between cuticle production and mechanical stress. If *k* refers to the substrate or film, the growth tensors for each subdomain are:

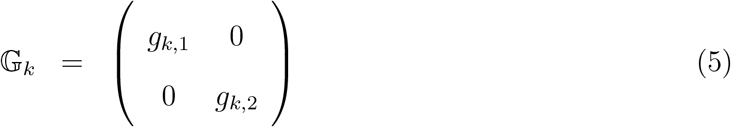

for the dynamics of cross-section only. Whereas, for the full volumetric case:

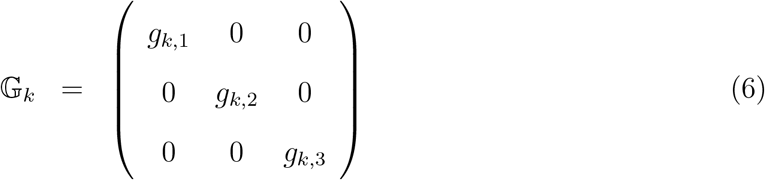

Any deformation that solves Eqn 4 is a result of cuticle growth and it is irreversible as there are no external forces acting on the volume. The source of compressive stress is therefore provided by the boundary conditions. The cuticle is synthesised locally on the surface of each cell and if the rates of expansion exceed those of cell in-plane elongation in a given direction, then we are trying to accommodate a larger volume within a smaller one, due to the presence of the neighboring cells (Fig. 3), in a similar fashion as the expansion of a cake is constrained by the baking container.

### Film thickness and turgor bulging

We start by presenting results of planar simulations which emulate the deformations of a cross-section. This is convenient because all of the ingredients responsible for the pattern formation are present at a relatively inexpensive computational effort, allowing us to extend the study to more elaborate settings. The initial stress free configuration is shown in Fig 3 and its given by Ω_*o*_: [0, *l_o_*] × [0, *H_o_*] with *H_o_* = *h* + *H_s_*. We choose the values *l_o_* = 2.0, *H_o_* = 0.2, *h/H_o_* = 0.008, *R* =10 and *ν* = 0.45, to mimic the aspect ratios observed in the cryo-fractures shown in Figs 1 and 2.

In Fig 4 we show results of two different sets of growth tensors for the film and for each case two different boundary conditions, the substrate in these cases does not expand. The growth tensors used are shown in the figure. The ones labeled *A* (left column) are given by *g_j,f_*(*t*) = 1 + *C_j_t*^2^, with *C*_1_ = 1 and *C*_2_ = 10*C*_1_ and *t* ∈ [0, 1], for the ones in *B* (right column) the expansion laws are the same but the thickness stops increasing at *t* = 0.5 before reaching the threshold and while the width of the film keeps increasing. This is done to investigate the effect of the thickness at the moment of reaching the buckling threshold and afterwards. The growth rates are relative to the reference configuration and the principal directions of volume growth (see Fig 3).

**Figure 4:**
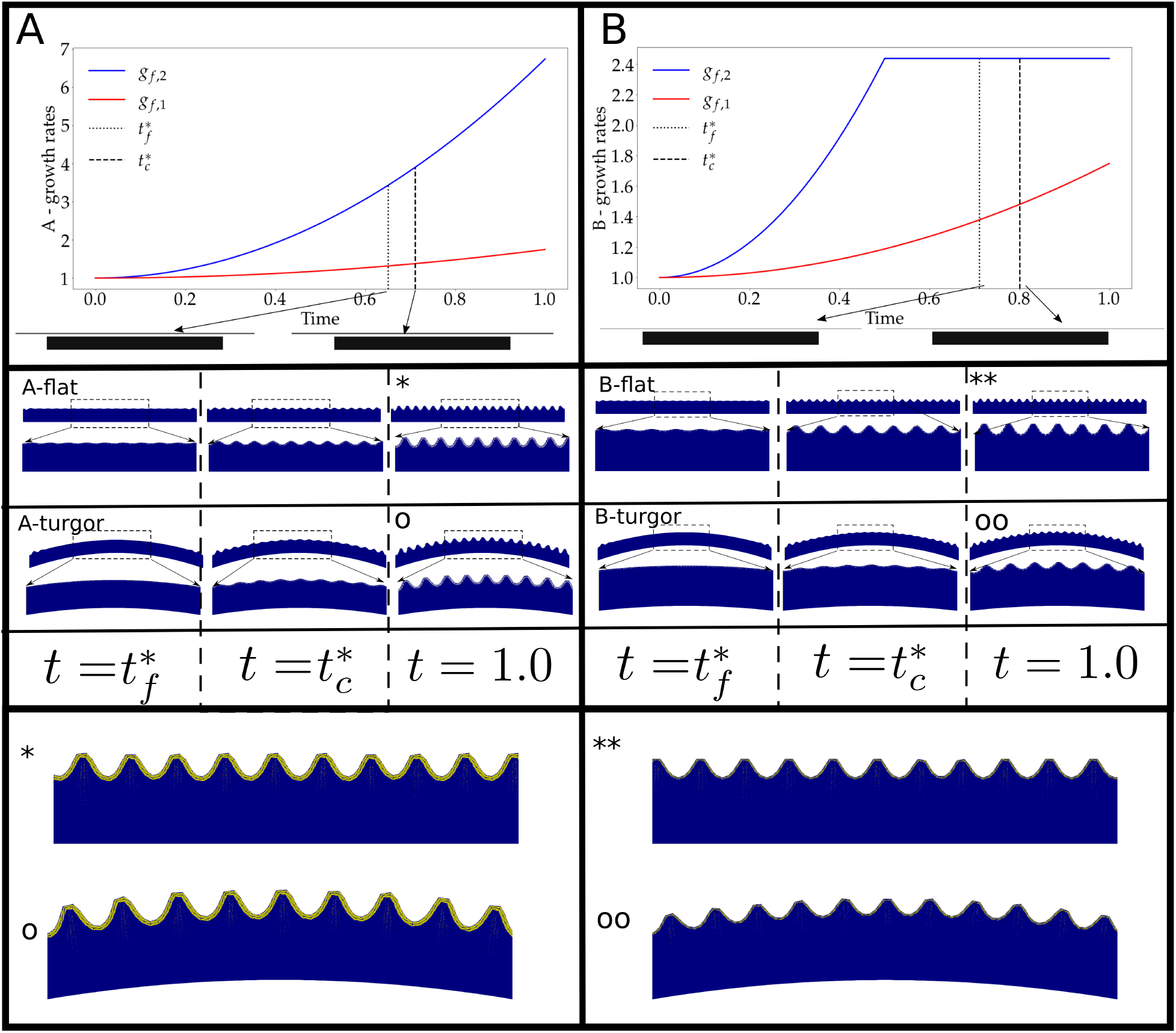
Effect of curved boundaries on triggering the instability. Solutions of the model using two sets of film expansion rates and boundary conditions. Column (A) Both entries use *g_f,k_*(*t*) = 1 + (*C_k_*/2)*t*^2^, with *C*_1_ = 1.0 and *C*_2_ = 10. Column (B) is the same as in (A) but with *g*_*f*,2_ stopping at *t* = 0.5. Each case is shown using flat and curved bottom boundaries. The bottom panel of each column shows a close-up of the solutions at *t* = 1.0 labeled by the symbols *, **, o, oo accordingly. For both growth scenarios the instability threshold is reached before using a flat boundary condition than for the curved cases. The reference configuration is the same for all cases, and the bottom boundary is updated in the same way as the growth tensor entries using a function *u*(*x*) = *r*(*t*)*x*(*x* – *w_o_*) with *r* = 0.2*t* for *t* < 0.65, this is done to emulate the bulging of cells due to turgor pressure as observed in Figs. 1 and 2.

For each set of growth tensors, the two boundary conditions used correspond to a case in which the bottom of the system remains flat and a case in which the bottom boundary is given by *f*(*t*) = *r*(*t*)*x*(*x* – *w_o_*), with *r*(*t*) = *kt*, *k* = 0.2 for *t* < *τ* = 0.65 and *r*(*t*) = *kτ* otherwise, which emulates bulging due to turgor pressure. These results are also shown in Fig. 4 and are included to illustrate several important concepts and effects.

“Time” has a specific meaning in the model. The solutions of (4) are obtained iteratively as elastic responses to updates of *g_f,k_* and *r*. Therefore, *t* controls relative changes due to growth for two or more internal parameters simultaneously. However, these solutions correspond to stationary configurations ignoring any transient dynamics, and in this respect time can be regarded as an internal clock-type parameter.

Regardless of the thickness growth, if the base of the substrate is deformed by increasing *r* then more expansive growth *g*_1,*f*_ is necessary to reach the stress buckling threshold. Here, the deformation of the base induces a strain across the entire domain, particularly in the cuticle film layer. Such stress needs to be compensated for in order to reach the buckling threshold. An immediate consequence of the delay is that, if the thickness of the film increases during that period, then the pattern wavelength is expected to be larger in accordance to (1). For the case in which the thickness stops growing before reaching the threshold, the delay also occurs but the change in wavelength is far less dramatic, the small difference is due to the extra length induced by the curved boundary, which allows the accommodation of an extra bump. Fig. 4 shows the resulting patterns for each of the growth tensors and boundary conditions at the onset of the flat systems 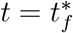, at the onset of the curved cases 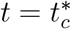 and at the end of the computation *t* =1.

Once the pattern has been triggered, the effect of further film expansion in the first principal direction is to increase the amplitude of the pattern without changing the wave-number. The morphology remains stable at least until a secondary instability is triggered [26, 27, 23, 28]. Here we focus only on the primary instability as it is the main pattern observed in *H. trionum*.

Fig. 5 shows the amplitudes and wavelengths of the solutions shown in Fig. 4 alongside measurements of the same properties from *H. trionum* petals at several stages of development. From the figure we see that for both sets of growth tensors, we obtain amplitudes and wavelengths in good agreement with those of the flowers by setting the width of the domain *w_o_* =< *w* >_12_, where *w_o_* is the initial width of the cell. However, some effects are of note. First the buckling threshold in the case in which the film thickness does not stop growing is slightly lower than the case in which the thickness expansion is stopped. This is expected as, in general, points in the deformed, current state are linear combinations of the local principal directions, therefore contain contributions of both growth laws (Figure 3). Second, in accordance to Eqn. 1, the case in which the threshold is reached with a thicker film selects a pattern with a larger average wavelength.

**Figure 5:**
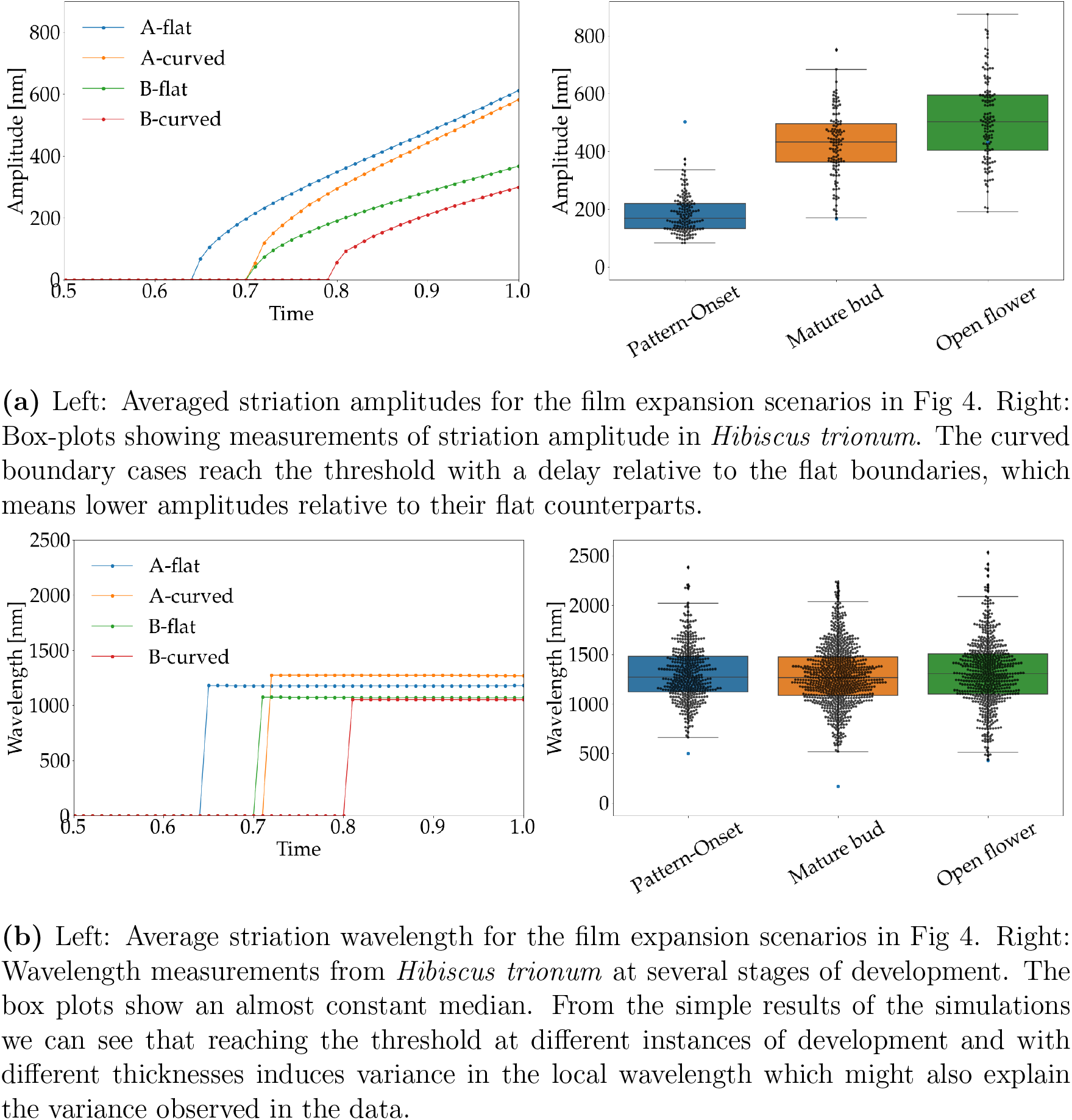
Amplitude and wavelength changes during *Hibiscus trionum* petal development and the effect of curved base boundaries. (a) The surface instability of the layer is triggered once a critical growth threshold is reached. The effect of a continuing film expansion after the pattern is triggered is the increase of the amplitudes. The pattern takes longer to fully populate the surface on curved substrates compared to the flat case. This implies that different curvature will lead to different amplitudes, which contributes to the variance observed in the data shown. (b) The wavelength across the surface as the amplitude increases remains constant in flat domains, which is not true in curved substrates. The average wavelength across the domains for curved and flat domains also a non-zero difference, between all the cases considered here.

Comparison of the cases using the same growth tensors but with different boundary conditions, namely, flat substrate base and curved surface base, reveals a delay in reaching the stress threshold to trigger the instability in curved substrates. This provides a good indicator of why the pattern is triggered in patches mostly on top of the junctions between cells, and as shown in Fig. 2b. where it is shown that the pattern populates flatter domains first then regions with larger local curvature.

### Multiple cells and substrate expansion

In order to explore in more detail the effects of epidermal cells with slightly different widths and cuticle base deformations, we now present results using a boundary condition which emulates the turgor induced curvature. We achieved this by dividing the bottom boundary of a reference configuration into three main segments of lengths 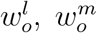 and 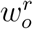 located between small clamped intervals. Each of the segments is subject to the conditions 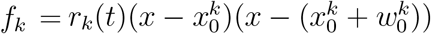 where *k* ∈ {*r,m,l*} stand for right, middle and left, and 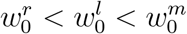.

Using the growth tensors provided in Fig. 6b we emulate the pattern development over cells of different width and turgor induced curvature. Solutions at the onset and fully formed patterns are shown in Fig. 6a, for the scenarios labeled *A* to *E*.

**Figure 6:**
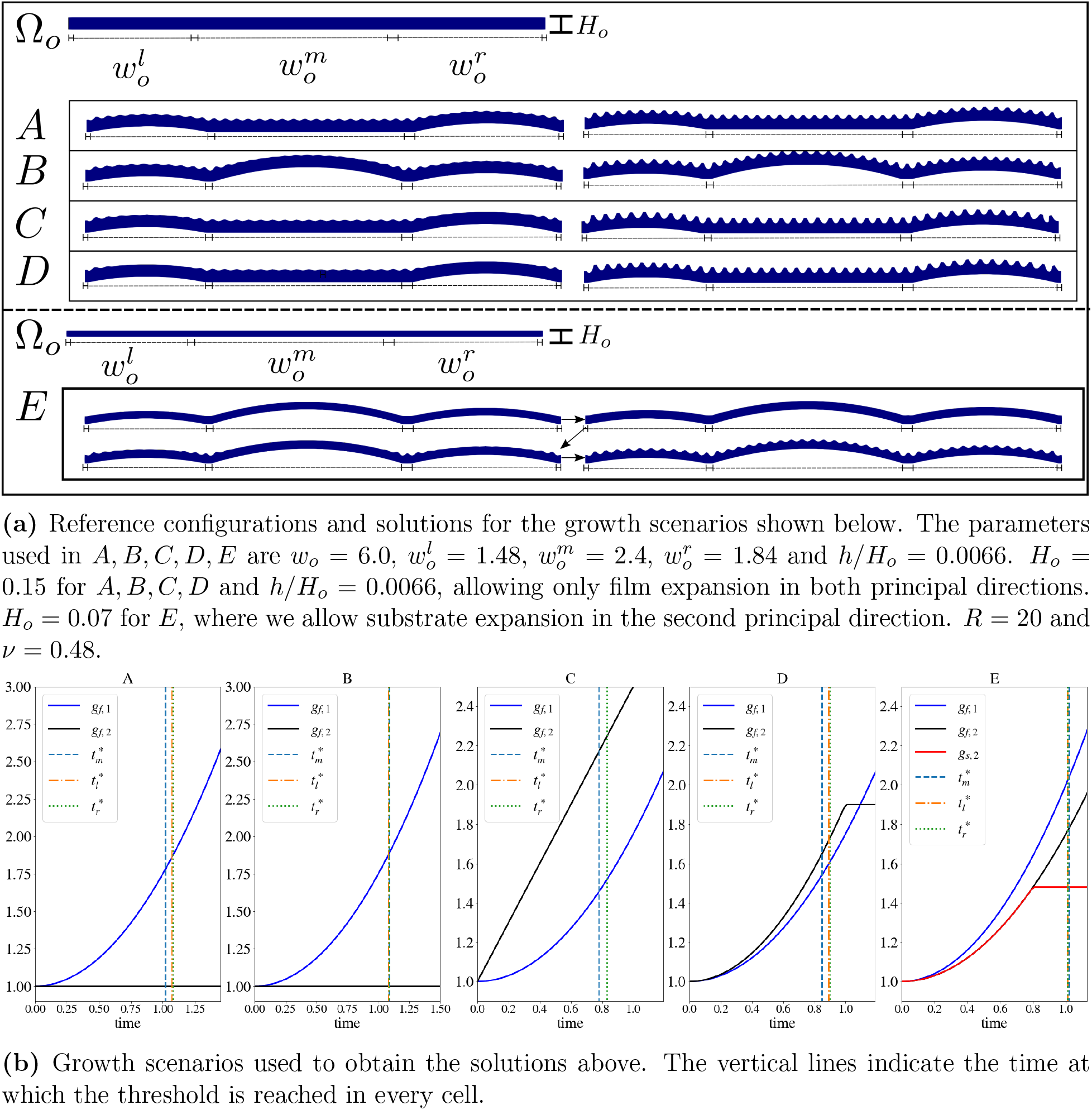
Multiple cells and non-uniform base deformations. (a) Solutions obtained from emulating three cells of slightly different length and growth rates as shown in (b). From this figure we can see that the process of pattern formation is well described by the model. The pattern starts forming in regions above the cell junctions and in flat regions. The solutions (A-D) shown show the configuration in which the pattern has populated the region for the first time and the final configuration. The sequence in E corresponds to the case in which the substrate thickess is also allowed to increase, the pattern formation process however still follows the same steps. The average wavelength and amplitudes for these solutions are shown in Fig 7.

Cases *A* and *B* are subject to the same growth rates, allowing only film expansion in the first principal direction, with *r_m_*(*t*) = 0 for *A*. In *B* every sub-domain is curved and given by the same rule as for the curved cases in Fig. 4. For *C* and *D* the film expands in the two principal directions, keeping the middle domain flat.

In case *E* we also consider growth of the substrate. Again, the buckling threshold is reached earlier in those sub-domains which remain flat (*A*, *C*, *D*) and the segments for which the boundary corresponds to the clamped junctions. A consequence of the delay in reaching the threshold is that the wavelength across the domain becomes less regular than for the flat case. This effect is partially geometrical and partially elastic. The geometrical effect is easy to understand if one considers a periodic flat curve described by (*x*, *h*(*x*)) and maps it into a curved convex segment given by **g** = (*x*, *g*(*x*)). The parametrisation 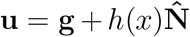 achieves this, where 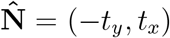, *t_x_* and *t_y_* are **g**’s unit normal and the components of its tangent field, respectively.

If *h*(*x*) is for instance *A cos*(*kx*), we can map the values at 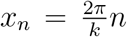 to points **u**_*n*_ = (*x_n_* – *At_y,n_,g_n_* + *At_x,n_*) and compute 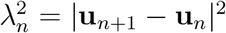, which can be written as:

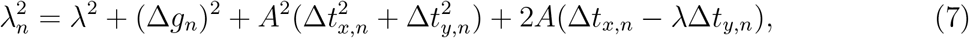

With 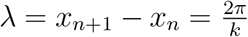, Δ*g_n_* = *g*_*n*+1_ – *g_n_*, and Δ*t_j,n_* = *t*_*j,n*+1_ – *t_j,n_*.

The first two terms in the right hand side of Eqn 7 are the hypotenuse of a triangle spanned by the difference in height in the concave profile and *h* evaluated at consecutive maxima. The last two terms make explicit that the larger the amplitude, the larger the separation between two consecutive peaks as these radiate away due to the concavity of *g*(*x*). This latter contribution is weaker as it only depends on differences in tangents evaluated at consecutive *x_n_*. This simple observation shows that the wavelengths across the concave curve are not constant, which explains in part the large variance observed in *H.trionum* petals (Fig. 5b). Another source of variance is set by mechanics, once the threshold is reached in a given sub-domain with a given wave-number, then, the amplitudes start to increase asynchronously with respect to other sub-domains with a different local geometry, inducing relative differences in the wavelengths.

Average values of each sub-domain and across the full domain of amplitudes and wave-lengths are shown in Fig. 7 for each case shown in Fig. 6. For *A* and *B*, where the only difference is the boundary condition of the middle cell, the average wavelength across the full domain is very similar, although due to the delay in reaching the threshold across the whole domain it takes more time to reach a similar set of amplitude values. For the growth scenarios *C* and *D* the film is allowed to expand anisotropically in the two principal directions. As expected the effect of reaching the buckling threshold with a larger thickness leads to an increase in the pattern average wavelengths.

**Figure 7:**
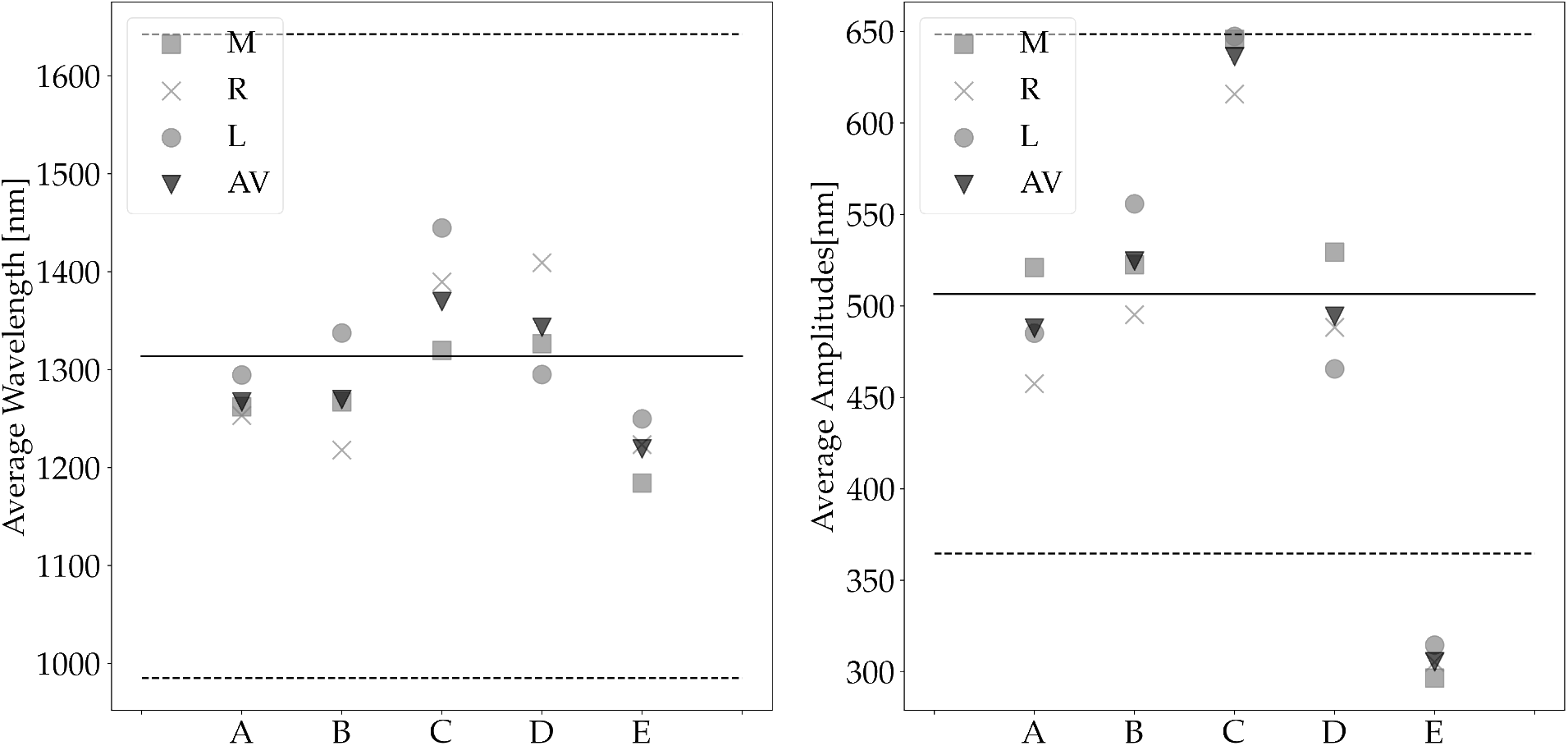
Average wavelength and average amplitude for each scenario in (a). These are scaled using the same factor as for the single cell case. The mean values and the standard deviation obtained from measurements of those quantities in *Hibiscus trionum* are also shown (solid and dashed lines). The labels *M, R, L* stand for middle, right and left cuticles whereas AV is the average value across the domain.

In all four cases the non-uniform turgor boundary conditions induce differences in the required film expansion to reach the threshold locally. If the film also increases its thickness, there is a contribution which might reduce the expansion required to buckle. In general, however, curved boundaries always require larger in-plane expansion with respect to the flat case to buckle. The non-homogeneous boundaries induce wavelengths which exhibit variations across each sub-domain but on average are still qualitatively described by a scaling law like Eqn. (1).

Case *E* corresponds to a case in which the substrate increases its thickness by setting *g*_*s*,2_ = *g*_*f*,2_ for *t* ∈(0, 0.8) after which the deposition process stops. The film continues its expansion in both principal directions as indicated in the figure. The pattern formation follows a similar process as those described in the previous cases. This suggests that, to an extent, the thickness expansion of the substrate in the process of buckling does not play a relevant role on its own. To investigate the role of substrate growth further, we briefly present the results of studying the system in a full three dimensional setting.

### Volumetric case

First, we present results of considering a reference configuration Ω_*o*_ = [0, *w_o_*] × [0, *l_o_*] × [0, *H_o_*] with *l_o_* = *w_o_* = *H_o_*/10, *H_o_* = 1 and *h* = *H_o_*/41 as shown in panel A of Fig. 8. The solutions are obtained imposing the boundary condition (*x, ty/l_o_*, 0.1*t* sin(*πx/w_o_*) sin(*πy/l_o_*)) on the base boundary surface to emulate turgor deformation. This boundary is stretched alongide the lateral surfaces in one preferred direction, emulating the expansion of the underlying cell. This is done by mapping the surfaces (*x, y*, 0) into (*x, ky*, 0), (0, *y, z*) into (0, *ky, z*) and (*w_o_,y,z*) into (*w_o_,ky,z*). The surfaces at the top and bottom of the length direction are free. The growth tensor entries are updated as: *g*_*f*,1_ = *g*_*f*, 2_ = *g*_*s*, 3_ = 1 + *t*. The amplitude of the turgor conditions is given by 0.1*t* and *g*_1,*s*_ = *g*_2,*s*_ are also set to be 1 + *t* but only for *t* < 0.2.

**Figure 8:**
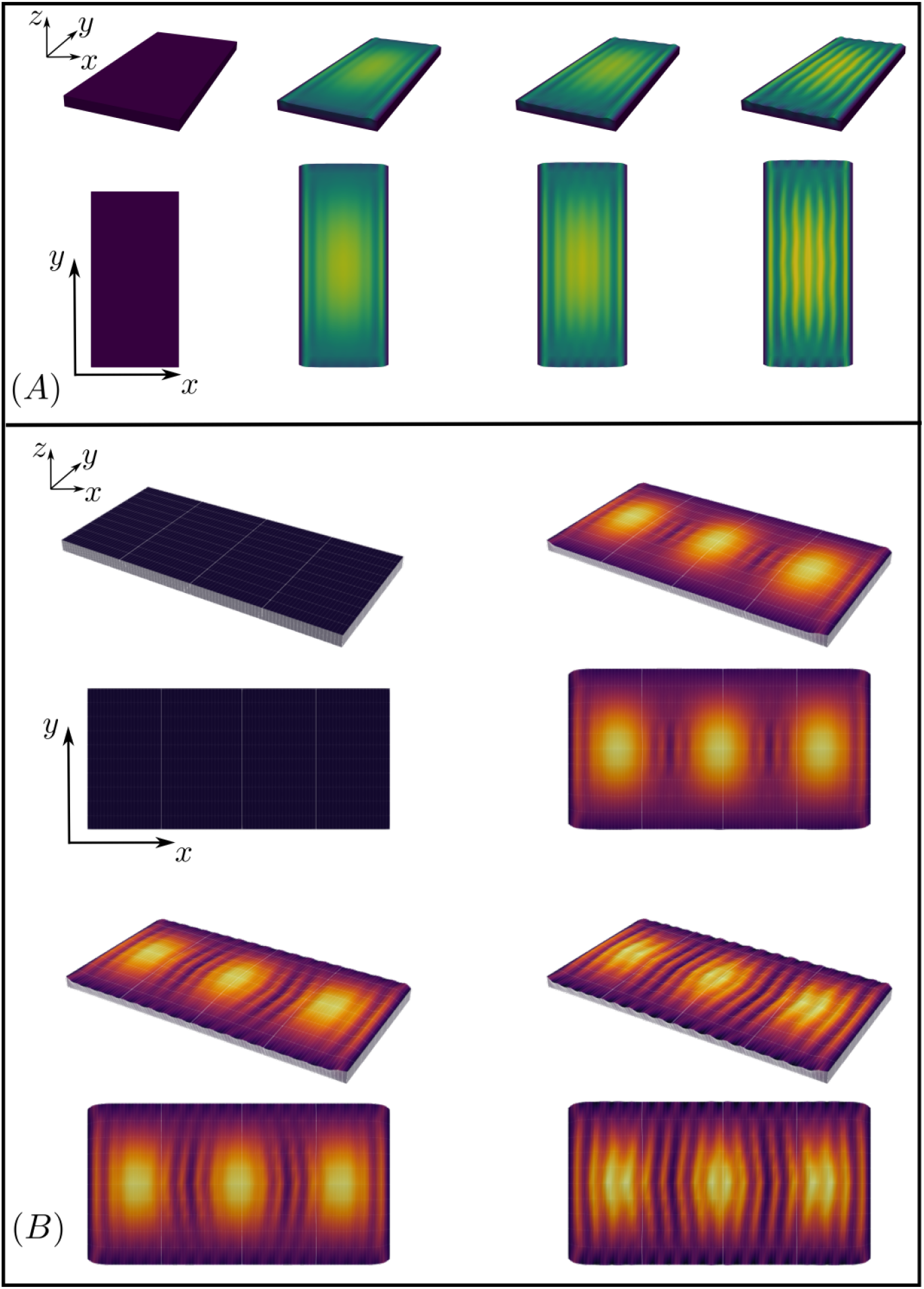
Volumetric growth. (A) Full volumetric growth, except for the film thickness. The base of the volume emulates deformation due to turgor pressure. The initial asymmetry and elongation overcome the growth induced stress in the elongation direction. (B) Solutions obtained using a bottom surface boundary condition which emulates three cells. The pattern is triggered in the flatter regions and the cell junctions, which adds some disorder to the final configuration.

The turgor deformation effect is the same as for the planar simulations, namely a delay in reaching the buckling threshold in those points with larger bulging. The asymmetric initial configuration and the subsequent elongation of the domain in the length direction suffices to develop the oriented pattern, despite the in-plane isotropic growth. A slight undulation due to the growth induced stress in the elongation direction can be seen, however it becomes less noticeable as the volume keeps elongating. This is also observed in the flowers during development, in the mature bud stage we notice slight undulations in the striations (Fig. 1 J) that disappear in the open flower once the cell length increases further (Fig. 1 K).

The addition of volume into the bulk of the substrate and the film are necessary to describe the observations of the cuticle as the petal develops, as stretching would result in a decrease of the thickness, which is not observed.

Finally, panel B in Fig 8 shows a similar numerical experiment to the case of three cells discussed earlier. In this case a region Ω_*o*_ = [0, *l_o_*] × [0, *l_o_*] × [0, *H_o_*] is used as reference configuration and the bottom boundary given by *u*(*x,y,t*) = *r*(*t*) sin^2^(*k*_1_*x*) sin^2^(*k*_2_*y*), with *k*_1_ = 3*π/w_o_* and *k*_2_ = *π/w_o_*. This emulates three underlying cell bulges in the domain. The remaining boundary conditions are the same as in the previous case.

The sequence shows results analogous to the three cells planar results. Specifically, the pattern is initiated at the regions which are flatter and the junctional regions. However, this example illustrates that the asynchronous pattern onset leads to configurations which are less ordered than the planar case. Again the source of in-plane stress is due to the lateral boundary conditions. The period and functions over which the turgor amplitude increases and these in-plane substrates grow are given by *r*(*t*) = *at*, *g*_1,*s*_ = *g*_2,*s*_ = *bt* + 1, for *t* < *τ* and *t* < 2*τ* respectively (*τ* = 0.12), whereas the film changing entries are *g*_1,*f*_ = *g*_2,*f*_ = 1 + *bt*. The growth rate of the domain boundaries in the length direction is *y*(*t*) = 0.5 * *bt*. All these with *a* = 0.2, *b* = 1.5 and *t* ∈ [0.0, 0.25].

The figure shows that the pattern starts forming in the region between the bulges where the maxima of *u* are located (sin^2^(*k*_2_*y*) = 1) and in the vicinities of these peaks, then the pattern populates those regions with flatter support, and finally it populates the domes. This asynchronous pattern triggering coincides with our observations of pattern initiation in *Hibiscus trionum*. The prescence of three domes also generates a pattern which appears more ‘organic’ than the previously discussed single cell case. In the real system, these effects are of course enhanced by different curvatures, different cell sizes and divisions of the underlying cells as well as irregularities in the cuticle material properties and geometric features.

## Discussion and Conclusions

By tracking the development of the cuticular striation pattern on petals of *H. trionum*, we have provided a clear picture of how such patterns are triggered. As the cells elongate in one preferred direction and the area of the petal increases, the in-plane area of the surface cells increases, as does the thickness of the top layer of the cuticle, the cuticle film. The increasing area of the average cell surface leads us to postulate expansion of the cuticle as the main mechanism behind the compressive stress responsible for the surface buckling. This is compatible with previous suggestions of ‘cuticle overproduction’ as the driving force. Equipped with this assumption, we then present a class of models which incorporate volume expansion into mechanical responses of a bilayer system.

In our models, cuticle production is interpreted as a local volume expansion of the layers, which results in in-plane stress. If the stress produced reaches a threshold then a pattern is triggered (critical growth condition), the threshold is determined by the stiffness mismatch ratio *R* > 1 condition.

A first thing to notice is that buckling instabilities are triggered only after in-plane stress in a given direction exceeds a threshold value. In our case, this translates into the critical growth condition. If the expansion of the layer does not occur, then the system under consideration will not develop a buckled surface pattern. Also, even if cuticle is continuously produced, if the relative stiffness of the two layers is close to *R* ≃ 1 then a stable pattern will not form.

Therefore, a mono-layer system or a bilayer far from the threshold are adequate models for cuticles which retain a flat surface.

The scenarios explored here generate buckled morphologies following developmental paths resembling those of the *H. trionum* petal (see Figs 2, 5, 7 and 8). Specifically, by imposing non-homogeneous boundary sub-domains, patterns are triggered locally in those regions with smaller curvatures before populating the full domain (arrows in Fig. 2b).

Once pattern formation is initiated, the effect of continuing cuticle expansion is an increase in the pattern amplitudes, while the wavelength remains constant on average. However, local differences in curvature introduce differences in the amount of in-plane growth required to reach the threshold. This effect, in turn, causes the pattern to be triggered asynchronously, occuring first in flatter regions and at cell junctions. This explains why the pattern is observed to develop in patchy domains, before covering the fully ridged petal sub-domain in *H. trionum*. The delay in reaching the buckling threshold has been numerically and analytically studied in [29] for systems with a curvature field present in the reference configuration. The asynchronous onset of the pattern and variations in the curvature over each cell induce variability in the wavelength values. We have provided a simple expression that accounts for the variability observed in the data obtained as a result of purely geometrical considerations (Eqn. 7), without appealing to differences in growth rates and/or spatial variations of material properties.

The results presented here are successful in describing the formation of the patterns present in *Hibiscus trionum* as a passive response to growth of bilayers constrained by boundaries, provided that *R* > 1. However, how these two conditions are met in some petal domains and not in others is still an open question, which requires a better understanding of the processes involved in cuticle formation and its material properties [25, 9, 30]. It is often assumed that new polymer chains are added exterior to the cell wall independently of the cell elongation process [31] as the result of active transport processes. However, it is pertinent to ask whether cell expansion plays a role in the fulfilment (or not) of the condition *R* > 1, as cell elongation requires regulated dynamic cell wall softening. How the two layers (film and substrate) differentiate and how cuticle production is coupled with elongation [32, 33] are questions which require an understanding of the regulation of cuticle development.

The morphoelastic approach taken here does not take into account any viscoelastic effects, which play an important role in the development of certain plant aerial parts. For instance other studies [34, 8, 7] have shown that tomato cuticles should be described as viscoelastic materials. Hyperelastic materials, such as the Neo-hookean material used here, are an excellent proxy for investigation of viscoelastic systems, provided the timescales of the material stress relaxation are slow with respect to the stresses acting upon the material [35].

The present work focuses on the primary instability as observed in *Hibiscus trionum*, and its role as the cause of structural colour in the petal. However the present class of models can be used to study the appearance of further instabilities which might be present over different systems as shown in [11]. The findings in our paper contribute to the broader literature on buckling instabilities in mutilayered systems, which have been used to study periodic mechanical patterns in epithelial layers [36, 37] and organ development [38, 39, 40]. Our work represents a comprehensive account of the mechanical ingredients for nano-scale pattern formation in the cuticle of petal epidermal cells.

## Materials and Methods

### *Hibiscus trionum* geometric measurements

- Plant growth conditions: *Hibiscus trionum* seeds were obtained from Cambridge University Botanic Garden. Plants were grown in a greenhouse with a controlled temperature of 21°C, and a 16 hour day light regime. Plants were grown using Levington’s M3 compost.
- Microscopy: Cell measurements were performed using a stereo microscope M205 FA-Leica. Film thickness was calculated using Cryo SEM fractures taken with a Zeiss-Quorum cryoSEM in the Sainsbury Laboratory Microscopy Core Facility with 3nm coating. We measured the wavelength using a WHX-5000 Keyence with a VH-Z500R/W/T objective and a 2000X magnification.

### Numerical solutions

To solve Eqn. 4 we implemented a Lagrangian description on a finite element scheme using the FEniCS framework [41], with dolfin version 2019.1.0. The problem is solved in variational form by specifying the weak form:

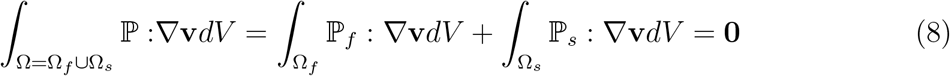

Where 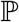 is obtained using the strain energy densities of a compressible Neo-Hookean material model:

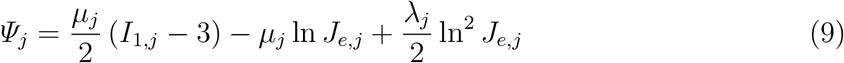

Where 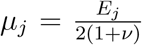 and 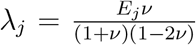 are the Lamé material parameters expressed in terms of *E_j_* and *ν*. Which for materials with the same *ν* satisfy: 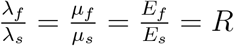.

The elastic deformation gradients 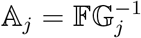 and the Cauchy-Green tensors 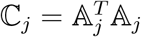 allow us to compute, *I*_1,*j*_ and *J_e,j_* and *J_g,j_*, the first and third principal invariants of 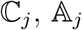 and 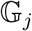 respectively. Alongside the second Piola-Kirchoff tensors:

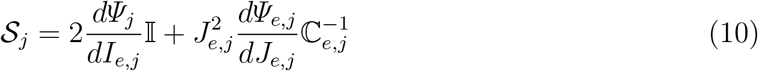

and:

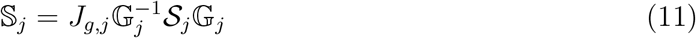

Which gives the desired stress tensors:

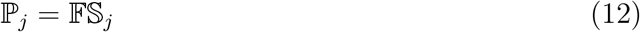

For the results presented here, we iteratively solve the system Eq. 8 in an interval *t* ∈ [0, *t_f_*] by updating the growth tensors at every time-step according to some growth laws, as described in the main text. In this respect, the time dependence is a quasi-static process as we compute stationary only configurations after an increase in volume relative to the reference configuration Ω_*o*_ as sketched in Fig. 3.

Full codes and documentation for the results presented here and some other wrinkling systems can be found in [42]

## Author Contributions

C.A.L. research conception and design, data analysis, mechanical modeling and simulation, manuscript drafting, edition and revision, figure preparation; C.A.A. research conception and design, plant growth, data acquisition, microscopy, manuscript edition and revision, figure preparation; C.C. manuscript edition and revision; A.J.C. and B.J.G. revision, edition and final version of manuscript approval, research funding and supervision.

## Acknowledgements

We are grateful to Matthew Dorling for excellent plant care; Raymond Wightman for help with Cryo-SEM; and Jordan Ferria.

This work has been supported by the Human Frontier Science Program (HFSP, No. RGP0019/2017 to B.J.G.) and the BBSRC (BB/P001157/1 to B.J.G.). We acknowledge the support of the SLCU Microscopy Core Facility, supported by the Gatsby Charitable Foundation.

## References

[1] Edwige Moyroud, Tobias Wenzel, Rox Middleton, Paula J. Rudall, Hannah Banks, Alison Reed, Greg Mellers, Patrick Killoran, M. Murphy Westwood, Ullrich Steiner, Silvia Vignolini, and Beverley J. Glover. Disorder in convergent floral nanostructures enhances signalling to bees. Nature, 550(7677):469–474, oct 2017.

[2] Lilan Hong, Joel Brown, Nicholas A. Segerson, Jocelyn K.C. Rose, and Adrienne H.K. Roeder. CUTIN SYNTHASE 2 maintains progressively developing cuticular ridges in arabidopsis sepals. Molecular Plant, 10(4):560–574, apr 2017.

[3] Chao Chen, Chiara A. Airoldi, Carlos A. Lugo, R. Konane Bay, Beverley J. Glover, and Alfred J. Crosby. Flower inspiration: Broad-angle structural color through tunable hierarchical wrinkles in thin film multilayers. Advanced Functional Materials, page 2006256, oct 2020.

[4] H. M. Whitney, M. Kolle, P. Andrew, L. Chittka, U. Steiner, and B. J. Glover. Floral iridescence, produced by diffractive optics, acts as a cue for animal pollinators. Science, 323(5910):130–133, jan 2009.

[5] Silvia Vignolini, Edwige Moyroud, Thomas Hingant, Hannah Banks, Paula J. Rudall, Ullrich Steiner, and Beverley J. Glover. The flower ofHibiscus trionumis both visibly and measurably iridescent. New Phytologist, 205(1):97–101, jul 2014.

[6] Udita Uday Ghosh, Sachin Nair, Anuja Das, Rabibrata Mukherjee, and Sunando Das-Gupta. Replicating and resolving wetting and adhesion characteristics of a rose petal. Colloids and Surfaces A: Physicochemical and Engineering Aspects, 561:9–17, jan 2019.

[7] Eva Domínguez, José Alejandro Heredia-Guerrero, and Antonio Heredia. The biophysical design of plant cuticles: an overview. New Phytologist, 189(4):938–949, nov 2010.

[8] Eva Domínguez, Jesús Cuartero, and Antonio Heredia. An overview on plant cuticle biomechanics. Plant Science, 181(2):77–84, aug 2011.

[9] Trevor H. Yeats and Jocelyn K.C. Rose. The formation and function of plant cuticles. Plant Physiology, 163(1):5–20, jul 2013.

[10] Chiara A. Airoldi, Carlos A. Lugo, Raymond Wightman, Beverley J. Glover, and Sarah Robinson. Mechanical buckling can pattern the light-diffracting cuticle of hibiscus trionum. Cell Reports, 36(11):109715, sep 2021.

[11] R. L. Antoniou Kourounioti, L. R. Band, J. A. Fozard, A. Hampstead, A. Lovrics, E. Moyroud, S. Vignolini, J. R. King, O. E. Jensen, and B. J. Glover. Buckling as an origin of ordered cuticular patterns in flower petals. Journal of The Royal Society Interface, 10(80):20120847–20120847, dec 2012.

[12] Xiao Huang, Yu Hai, and Wei-Hua Xie. Anisotropic cell growth-regulated surface micropatterns in flower petals. Theoretical and Applied Mechanics Letters, 7(3):169–174, may 2017.

[13] Yanping Cao and John W. Hutchinson. Wrinkling phenomena in neo-hookean film/substrate bilayers. Journal of Applied Mechanics, 79(3), apr 2012.

[14] Derek Breid and Alfred J. Crosby. Effect of stress state on wrinkle morphology. Soft Matter, 7(9):4490, 2011.

[15] E. Cerda and L. Mahadevan. Geometry and physics of wrinkling. Physical Review Letters, 90(7), feb 2003.

[16] Jan Genzer and Jan Groenewold. Soft matter with hard skin: From skin wrinkles to templating and material characterization. Soft Matter, 2(4):310, 2006.

[17] Jan Groenewold. Wrinkling of plates coupled with soft elastic media. Physica A: Statistical Mechanics and its Applications, 298(1-2):32–45, sep 2001.

[18] Bo Li, Yan-Ping Cao, Xi-Qiao Feng, and Huajian Gao. Mechanics of morphological instabilities and surface wrinkling in soft materials: a review. Soft Matter, 8(21):5728, 2012.

[19] Basile Audoly and Arezki Boudaoud. Buckling of a stiff film bound to a compliant substrate—part i:. Journal of the Mechanics and Physics of Solids, 56(7):2401–2421, jul 2008.

[20] S. Cai, D. Breid, A.J. Crosby, Z. Suo, and J.W. Hutchinson. Periodic patterns and energy states of buckled films on compliant substrates. Journal of the Mechanics and Physics of Solids, 59(5):1094–1114, may 2011.

[21] Hamza Alawiye, Ellen Kuhl, and Alain Goriely. Revisiting the wrinkling of elastic bilayers i: linear analysis. Philosophical Transactions of the Royal Society A: Mathematical, Physical and Engineering Sciences, 377(2144):20180076, mar 2019.

[22] Hamza Alawiye, Patrick E. Farrell, and Alain Goriely. Revisiting the wrinkling of elastic bilayers II: Post-bifurcation analysis. Journal of the Mechanics and Physics of Solids, 143:104053, oct 2020.

[23] M.A. Holland, B. Li, X.Q. Feng, and E. Kuhl. Instabilities of soft films on compliant substrates. Journal of the Mechanics and Physics of Solids, 98:350–365, jan 2017.

[24] Alain Goriely. The Mathematics and Mechanics of Biological Growth. Springer New York, 2017.

[25] Joanna Skrzydeł, Dorota Borowska-Wykręt, and Dorota Kwiatkowska. Structure, assembly and function of cuticle from mechanical perspective with special focus on perianth. International Journal of Molecular Sciences., 22(8):4160, apr 2021.

[26] Fabian Brau, Hugues Vandeparre, Abbas Sabbah, Christophe Poulard, Arezki Boudaoud, and Pascal Damman. Multiple-length-scale elastic instability mimics parametric resonance of nonlinear oscillators. Nature Physics, 7(1):56–60, oct 2010.

[27] Silvia Budday, Ellen Kuhl, and John W. Hutchinson. Period-doubling and period-tripling in growing bilayered systems. Philosophical Magazine, 95(28-30):3208–3224, feb 2015.

[28] Qiming Wang and Xuanhe Zhao. A three-dimensional phase diagram of growth-induced surface instabilities. Scientific Reports, 5(1), mar 2015.

[29] Fei Jia, Simon P. Pearce, and Alain Goriely. Curvature delays growth-induced wrinkling. Physical Review E, 98(3), sep 2018.

[30] Dariusz Stępiński, Maria Kwiatkowska, Agnieszka Wojtczak, Justyna Teresa Polit, Eva Domínguez, Antonio Heredia, and Katarzyna Popłońska. The role of cutinsomes in plant cuticle formation. Cells, 9(8):1778, jul 2020.

[31] Daniel J. Cosgrove. Plant cell wall extensibility: connecting plant cell growth with cell wall structure, mechanics, and the action of wall-modifying enzymes. Journal of Experimental Botany, 67(2):463–476, 11 2015.

[32] Christopher E. Jeffree. The fine structure of the plant cuticle. In Biology of the Plant Cuticle, pages 11–125. Blackwell Publishing Ltd, 2007.

[33] Mike Pollard, Fred Beisson, Yonghua Li, and John B. Ohlrogge. Building lipid barriers: biosynthesis of cutin and suberin. Trends in Plant Science, 13(5):236–246, may 2008.

[34] G. Lopez-Casado, A. J. Matas, E. Dominguez, J. Cuartero, and A. Heredia. Biomechanics of isolated tomato (solanum lycopersicum l.) fruit cuticles: the role of the cutin matrix and polysaccharides. Journal of Experimental Botany, 58(14):3875–3883, nov 2007.

[35] Marco Amabili. Nonlinear Mechanics of Shells and Plates in Composite, Soft and Biological Materials. Cambridge University Press, oct 2018.

[36] E. Hannezo, J. Prost, and J.-F. Joanny. Instabilities of monolayered epithelia: Shape and structure of villi and crypts. Physical Review Letters, 107(7), aug 2011.

[37] E. Hannezo, J. Prost, and J.-F. Joanny. Theory of epithelial sheet morphology in three dimensions. Proceedings of the National Academy of Sciences, 111(1):27–32, dec 2013.

[38] M. Ben Amar and F. Jia. Anisotropic growth shapes intestinal tissues during embryogenesis. Proceedings of the National Academy of Sciences, 110(26):10525–10530, jun 2013.

[39] O V Manyuhina, David Mayett, and J M Schwarz. Elastic instabilities in a layered cerebral cortex: a revised axonal tension model for cortex folding. New Journal of Physics, 16(12):123058, dec 2014.

[40] Silvia Budday, Paul Steinmann, and Ellen Kuhl. The role of mechanics during brain development. Journal of the Mechanics and Physics of Solids, 72:75–92, dec 2014.

[41] Martin Alnæs, Jan Blechta, Johan Hake, August Johansson, Benjamin Kehlet, Anders Logg, Chris Richardson, Johannes Ring, Marie E Rognes, and Garth N Wells. The fenics project version 1.5. Archive of Numerical Software, Vol 3, 2015.

[42] https://github.com/calugo/wrinkles.

